# Genetic control of cell layer interactions in plants via tissue mechanics

**DOI:** 10.1101/2023.02.14.527838

**Authors:** Robert Kelly-Bellow, Karen Lee, Richard Kennaway, Elaine Barclay, Annabel Whibley, Claire Bushell, Jamie Spooner, Man Yu, Paul Brett, Baldeep Kular, Shujing Cheng, Jinfang Chu, Ting Xu, Brendan Lane, James Fitzsimons, Yongbiao Xue, Richard Smith, Christopher D. Whitewoods, Enrico Coen

**Affiliations:** Department of Cell and Developmental Biology, John Innes Centre, Norwich Research Park; Colney Lane, Norwich, NR4 7UH, UK; Department of Biochemistry and Metabolism, John Innes Centre, Norwich Research Park; Colney Lane, Norwich, NR4 7UH, UK; National Centre for Plant Gene Research (Beijing), Institute of Genetics and Developmental Biology, Chinese Academy of Sciences; Beijing 100101, China; College of Advanced Agricultural Sciences, University of Chinese Academy of Sciences; Beijing 100039, China; State Key Laboratory of Plant Cell and Chromosome Engineering, Institute of Genetics and Developmental Biology, Chinese Academy of Sciences; Beijing 100101, China; Department of Computational and Systems Biology, John Innes Centre, Norwich Research Park; Colney Lane, Norwich, NR4 7UH, UK; Sainsbury Laboratory, University of Cambridge; Cambridge CB2 1LR, UK

## Abstract

Plant development depends on coordination of growth between different cell layers. Coordination may be mediated by molecular signalling or mechanical connectivity between cells, but evidence for genetic control via direct mechanics has been lacking. We show that a brassinosteroid-deficient dwarf mutant of the aquatic plant *Utricularia gibba* has twisted internal tissue, likely caused by a mechanical constraint from a slow-growing epidermis creating tissue stresses. This conclusion is supported by showing that inhibition of brassinosteroid action in an *Arabidopsis* mutant compromised for cell adhesion, enhances epidermal crack formation, an indicator of increased tissue tension. Thus, genes driving brassinosteroid synthesis can promote growth of internal tissue by reducing mechanical epidermal constraint, showing that tissue mechanics plays a key role in coordinating growth between cell layers.

**One-Sentence Summary:** Internal twists in a mutant carnivorous plant reveal how genes control growth via tissue mechanics.

## Main Text

Many multicellular organisms are formed from multiple cell layers, raising the question of how growth is coordinated between layers to produce an integrated final form. In plants, evidence from genetic chimeras, or from layer-specific modification of gene function, show that genes active in one layer can act non-autonomously to influence growth in other layers, leading to coordinated overall growth (1-4). Non-autonomy could be explained through chemical signalling between layers, or by mechanical connectivity. The potential role of mechanics is illustrated by the phenomenon of tissue tension, in which the epidermis constrains the growth of internal layers, leading to bending and coiling when tissue is cut (5-8). However, little is known about how tissue tension is controlled genetically and thus the role it may play in non-cell-autonomous gene action. Here we address this problem through the analysis of dwarf mutants in the aquatic plant *Utricularia gibba* and in *Arabidopsis thaliana*.

*U. gibba* is a carnivorous plant, which in its vegetative phase comprises a spiral apex which produces stolons bearing filiform leaves and traps (9) (Fig. 1A). To investigate the genetic control of *U. gibba* development, we carried out a mutagenesis screen. Obtaining large numbers of progeny from *U. gibba* proved difficult because of poor seed set and germination rates. Rather than seed mutagenesis, we therefore mutagenized small stolon explants of *U. gibba* with EMS and grew each on to flowering (see Methods for details). M1 seed was collected from 441 explants and gave M2 phenotypes including, one M2 family that contained 2 dwarf and 2 wild-type plants. Seed from one of the wild types, gave 37 wild type, 9 dwarf, and 3 extreme-dwarf plants (Fig. 1, A to C). These numbers were consistent with segregation of two recessive mutations: one giving the dwarf phenotype, and the other enhancing the dwarf phenotype.

**Fig. 1.**
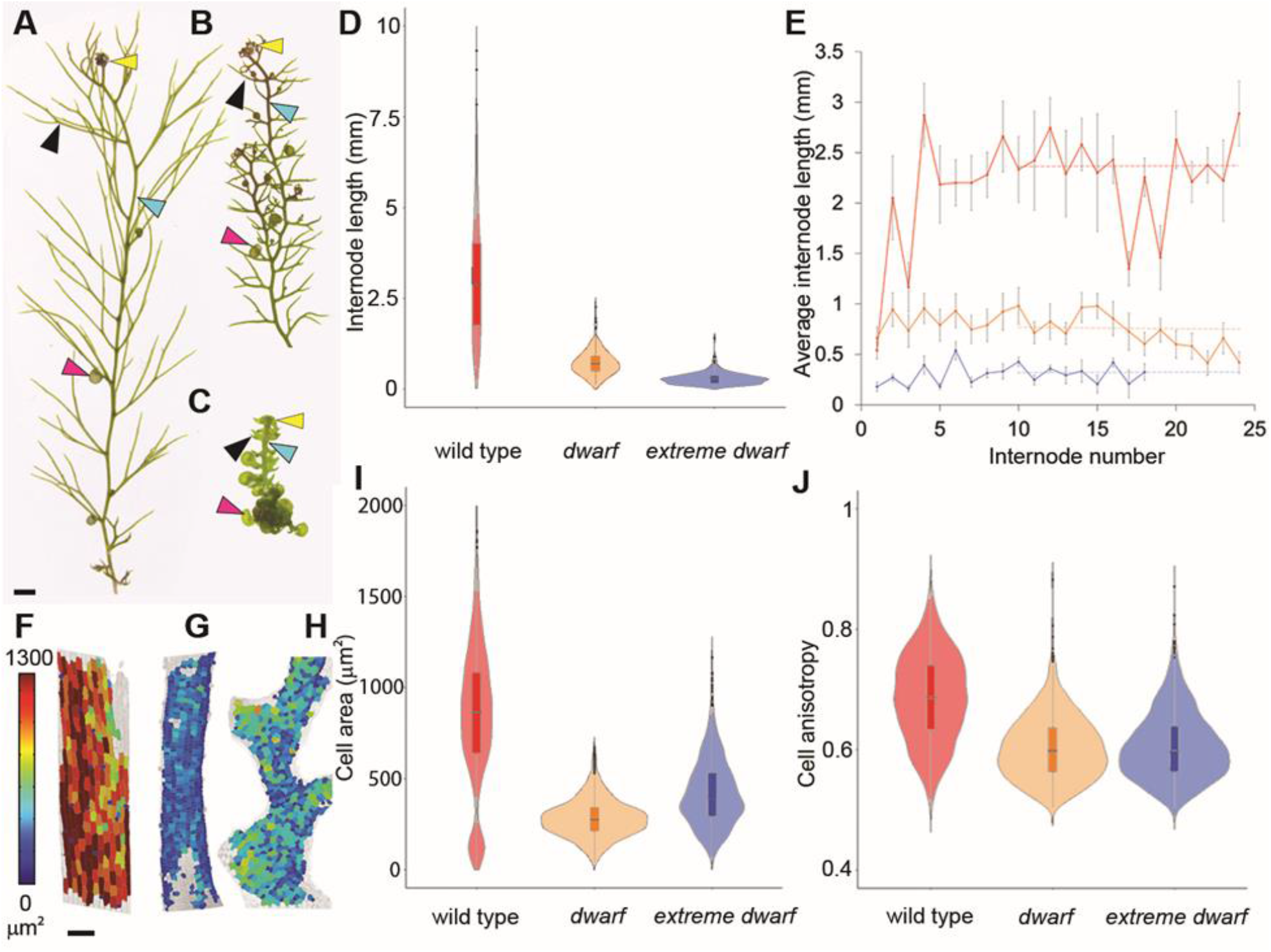
External phenotype of U. gibba wild type and dwarf mutants. (**A** to **C**) *U. gibba* vegetative plants comprise a spiral apex (yellow arrowhead), filiform leaves (black arrowhead), internodes (cyan arrowhead) and traps (magenta arrowhead). (A) Wild type. (B) Dwarf. (C) Extreme dwarf. Scale bar 1 mm. (**D**) Violin plots of wild-type (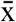 = 3.07 mm, n=10), dwarf (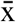 = 0.72 mm, n=10) and extreme dwarf (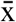 = 0.29 mm, n=13) mature internode lengths from plants grown in continuous culture. Block indicates interquartile range and horizontal line the mean. Both mutants have reduced lengths compared to wild type (*p* < 0.001). (**E**) Internode lengths from growing explants of wild type (red), dwarf (orange) and extreme dwarf (blue) plotted against internode number. Dashed line shows mean from internode 10 onwards. (**F** to **H**) Heat maps of cell area in mature stolons of wild type (F), dwarf (G) and extreme dwarf (H) mutants. Scale bar 100 µm. (**I** and **J**) Violin plots of cell area (I) and cell anisotropy (cell max length/(cell max length + cell min length)) (J) of mature stolons of wild type (n=1817 cells from 8 plants), dwarf (n=2289 cells from 5 plants) and extreme dwarf (n=1494 cells from 4 plants). Both mutants have significantly lower values than wild type (p< 0.001, I; p< 0.001 J).

Both the dwarf and extreme dwarf phenotypes were characterised by short internodes, short leaves and small traps (Fig. 1, A to D). To follow their development, internodes were numbered sequentially according to their position relative to the spiral apex, with the internode below the first clearly visible leaf denoted as 1 (fig. S1). Internodes of wild type increased in length until about internode 4 (Fig. 1E). Dwarf plants had a similar internode length to wild type at node 1 but exhibited very little growth after this. Extreme-dwarf internodes were shorter at node 1 than other genotypes and exhibited little subsequent growth.

Both epidermal cell area and anisotropy were reduced in mutants (Fig. 1, F to J). Measurements of cell length parallel to the stolon axis indicated that 70% of the reduction in dwarf plant internode length was caused by reduced growth after cell division arrest (fig. S2). Further reduction in internode length in extreme dwarf plants was caused by reduced growth prior to division arrest.

We next determined the phenotype of internal tissues. Wild-type stolons had a cylindrical epidermis connected by 5-6 straight “blades” to an axial vascular bundle, with air spaces between the blades (Fig. 2, A to C). Stolons of dwarf plants retained air spaces but had twisted blades (Fig. 2, D to F). Twisting extended to the internal vascular bundle, which was sinuous in longitudinal confocal sections rather than straight (Fig. 2D). Stolons of the extreme dwarf had smaller air spaces and less twisted vasculature than dwarf (Fig. 2, G to I).

**Fig. 2.**
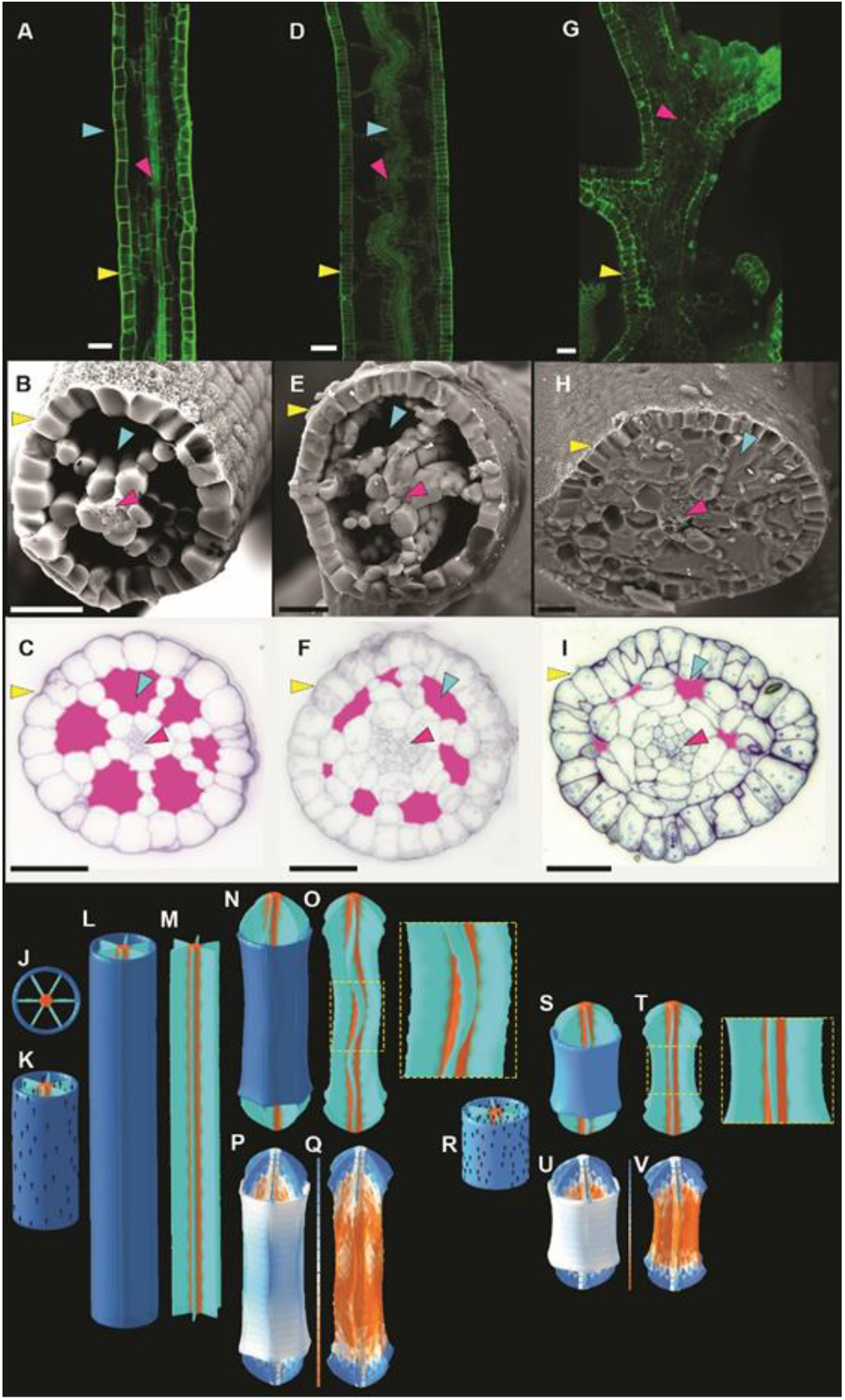
Internal phenotype and simulations of U. gibba wild type and dwarf mutants. (**A** to **I**) Wild type (A to C), dwarf (D to F) and extreme dwarf (G-I) longitudinal confocal sections (A, D and G), freeze-fracture scanning electron microscopy (B, E and H) and toluidine-blue-stained transverse sections (C, F, and I). Epidermis (yellow arrowhead), vasculature (magenta arrowhead) and air spaces (cyan arrowhead). Intercellular spaces highlighted with magenta. Scale bars 50 µm. (**J** to **V**) Computer simulations. (J) Top-down view of initial state showing epidermis (blue), blades (cyan) and axial core (orange). (K) Side view of initial state for wild-type and dwarf mutant models. Polarity shown with black arrows. (L) Final state when all tissue has the same specified growth rate (wild type). (M) as (L) with epidermis removed. (N) Final state when epidermis has a lower specified growth rate (dwarf). (O) as (N) with epidermis removed. (P and Q) as (N and O) showing tissue compression in orange and tension in blue. (R) Side view of initial state for extreme dwarf model. (S) Final state when epidermis has a lower specified growth rate (extreme dwarf). (T) as (S) with epidermis removed. (U and V) as (S and T) showing tissue compression in orange and tension in blue.

The twisted internal tissue of the dwarf plants might be caused by altered patterning of vascular and blade tissue, such that it differentiates in a contorted arrangement. Alternatively, twisting might arise through direct mechanical interactions between layers. To determine whether the mechanical interactions might be involved, we modelled growth of a hollow cylinder connected by blades to an axial core (Fig. 2, J and K). We distinguish between specified growth of a tissue region, which is how much the region would grow in mechanical isolation, and resultant growth, which is how much the region grows when mechanically connected to the rest of the tissue (10). Specified growth was oriented parallel to a polarity field running from the bottom to the top of the stolon (Fig. 2K). If all regions had the same specified growth rate, the initial cylinder elongated without any twisting of internal tissue (Fig. 2, L and M). However, if specified growth rate was much lower in the epidermis than the rest of the tissue, the growth conflict led to tissue tension in the epidermis and tissue compression in the blades and axial core, causing twisting as observed in the dwarf plants (Fig. 2, N to Q). If the cylinder was initially shorter (Fig. 2R), as observed for internodes of the extreme-dwarf (Fig. 1D), twisting was reduced (Fig. 2S to V), and further reduced if the air spaces were smaller (fig. S3), as also observed (Fig. 2I).

To distinguish between patterning and mechanical hypotheses, we determined the developmental timing of the twisted phenotype in dwarf plants. According to the patterning hypothesis, twisting should be observed as soon as vascular cell types can be distinguished. The mechanical hypothesis predicts twisting should occur after air-spaces have formed. Straight vasculature surrounded by air spaces was evident at internodes 0 and 1 of the dwarf mutant (Fig. 3A), as in wild type (fig. S4B). Twisted internal tissue in dwarf plants was not observed until internode 4 onwards (Fig. 3B). These findings support the mechanical hypothesis.

**Fig. 3.**
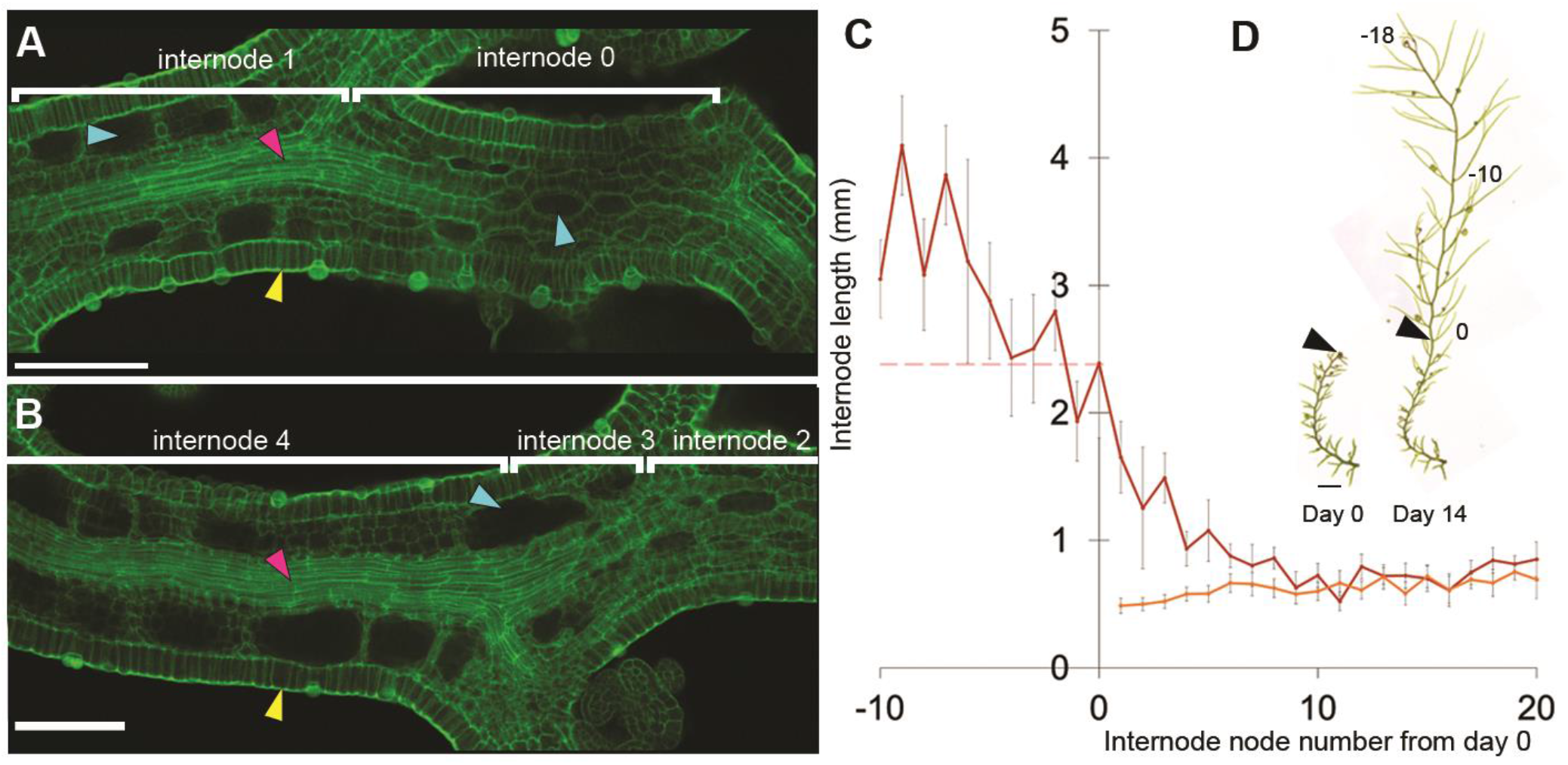
Early developmental stage of U. gibba dwarf and rescue by brassinosteroid treatment. (**A** and **B**) Confocal Z-slice of early dwarf internodes. Vasculature (magenta arrowhead), air spaces (cyan arrowhead), epidermis (yellow arrow). Scale bar 100 µm. (**C**) Average initial internode lengths of dwarf explants (day 0, solid orange line), and 14 days after treatment with 0.01 µM epiBL (day 14, solid brown line) (n≥ 10). Internodes were numbered according to their position on day 0, and corresponding internodes identified on day 14. Negative internode numbers were assigned by sequential counting based on the image at day 14. A significant difference (*p* < 0.05) between day 0 and day 14 internode lengths was observed for internodes 1 to 5, but not for higher internode numbers. Day 14 internode lengths were not significantly different from the mean wild-type mature internode length based on Fig. 1E (red dashed line) for internodes 0 to -10. Error bars show SEM. (**D**) Example of explant imaged on day 0 and day 14. Black arrowhead points to internode 0. Scale bar 5 mm.

To identify the *DWARF* gene, we sequenced the wild-type progenitor used for mutagenesis and 20 individuals from a segregating mutant family: 13 wild type, 5 dwarf and 2 extreme dwarf. Only one SNP satisfied the criteria for being at the *DWARF* locus: the SNP was absent from the progenitor, was heterozygous or absent in wild type segregants, and homozygous in all mutants. Genotyping a further 20 wild-type, 5 dwarf and 1 extreme-dwarf individuals confirmed this segregation pattern. The extreme-dwarf plants carried 4 additional SNPs absent from the progenitor that are candidate mutations in an enhancer of *DWARF* gene.

The *DWARF* SNP introduced an early stop codon in a gene encoding a cytochrome P450 90B1 enzyme, which catalyses the C22-alpha-hydroxylation step in the brassinosteroid (BR) biosynthesis pathway (11). This gene is homologous to *DWARF4* in *Arabidopsis*, which affects cell area, cell anisotropy in a similar way to *DWARF* (12, 13). Mass spectrometry on dwarf and extreme dwarf plants revealed absence of two BR precursors (castasterone and 6-Deoxocastasterone) after the C22-alpha-hydroxylation step (fig. S5), but presence of a precursor (campesterol) before the block (fig. S5). One BR precursor after the block (typhasterol) showed low levels in all genotypes which may reflect the coincident peaks (fig. S5). Growing *dwarf* mutants in 0.01 µM epibrassinolide (epiBL) rescued the *dwarf* phenotype (fig. S6). Extreme-dwarf was partially rescued for internode length (fig. S6B). Thus, *DWARF* likely encodes a BR biosynthesis gene.

To determine the timing of BR action on internode development, we tracked *dwarf* stolons after treatment with 0.01 µM epiBL. Internodes 0 to 5 showed a significant increase in average internode length (*p* < 0.05), compared to untreated controls, with treatments of internode 0 giving near-complete rescue (Fig. 3, C and D). Internodes more mature than internode 5 were not significantly affected by epiBL. Thus, BR likely acts during internodes 0 to 5 to promote epidermal growth and release internal tissue from mechanical constraint.

In contrast to the *dwarf* mutant of *Utricularia*, the *dwarf4* mutant of *Arabidopsis* does not give noticeably twisted internal tissue (12). BR may not affect tissue stresses in *Arabidopsis*, or BR may act as in *Utricularia* but lack of air spaces in *Arabidopsis* stems precludes internal tissue buckling (a solid cylinder model with reduced epidermal specified growth did give internal twisting (fig. S7)).

To distinguish these hypotheses, we exploited an *Arabidopsis* cell adhesion mutant, *quasimodo2-1* (*qua2-1*), which has weaker cell-cell adhesion and therefore provides an indicator of epidermal tissue tension through crack formation (14, 15). Consistent with previous findings, *qua2-1* hypocotyls had cracks that typically spanned 1 or 2 cell files (Fig 4, A, B and E). By contrast, *qua2-1* hypocotyls grown in the presence of a brassinosteroid inhibitor, BRZ, displayed significantly larger cracks (Fig. 4, C, D and E). Thus, inhibiting BR likely increases tissue tension in the *Arabidopsis* epidermis.

**Fig. 4.**
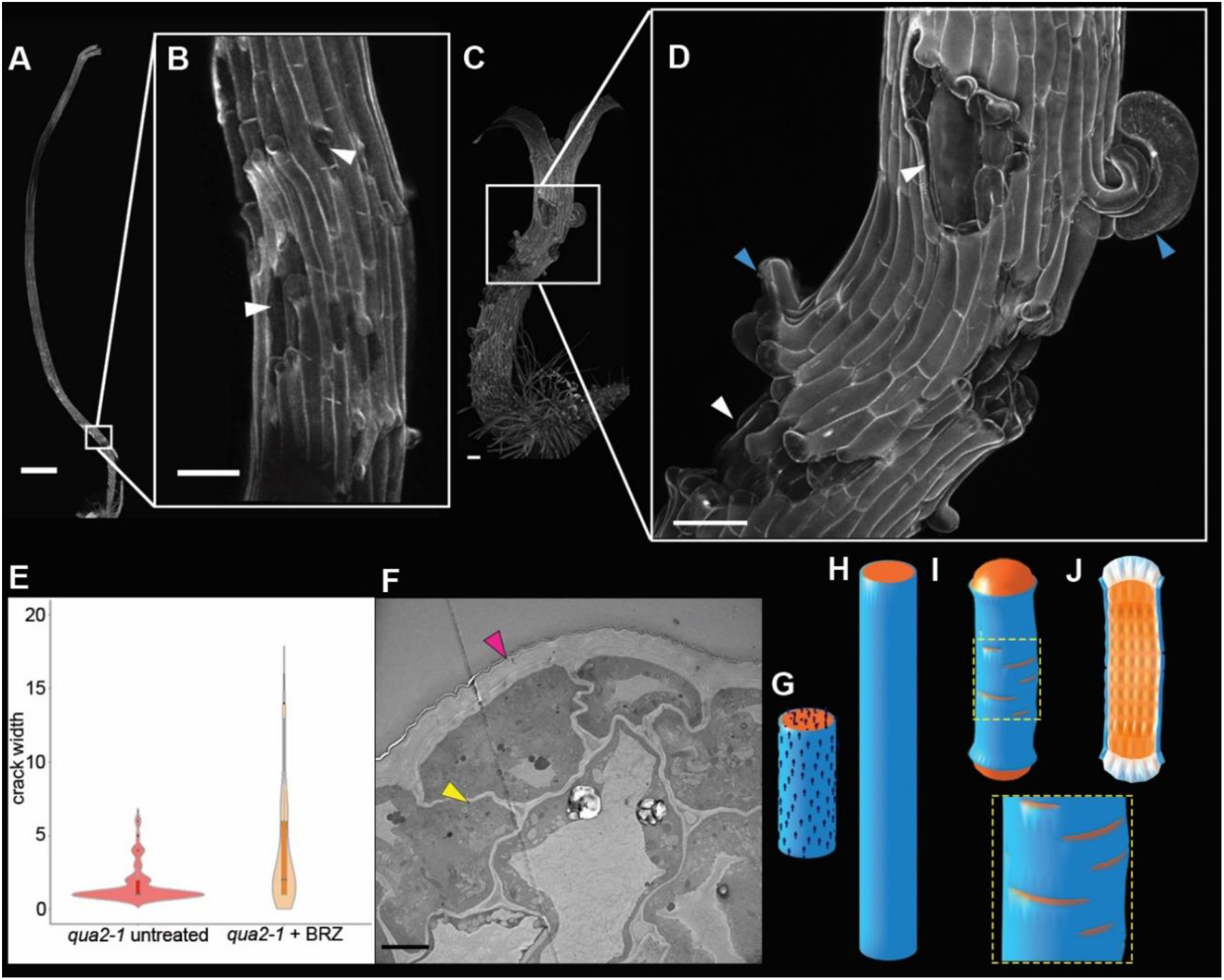
Phenotype and simulations *qua2-1* Arabidopsis hypocotyls treated with brassinosteroid inhibitor. (**A** to **D**) Confocal images of seedlings after 9 days growth in dark. (**A**) *qua2-1* (**B**) Close up of region highlighted in (A). (**C**) *qua2-1* grown on 1 µM brassinazole. (**D**) Close up of region highlighted in (C). White arrowheads indicate cracks, blue arrowheads indicate curved cells at crack boundaries. Scale bars 100 µm except (A) which is 1 mm. (**E**) Violin plots of crack widths. Mean crack width is significantly greater with brassinazole (BRZ) treatment (p < 0.001, untreated hypocotyls n = 2, n cracks = 74, BRZ-treated hypocotyls n = 6, n cracks = 83). (**F**) Transmission electron micrograph of wild-type hypocotyl epidermal cell after 4 days growth, showing thicker outer epidermal cell wall. Scale bar 5 µm. (**G** to **J**) Computer simulations of development. (G) Initial state, with outer epidermis in blue and inner regions in red. Arrows indicate polarity. (H) Final state with all regions having the same specified growth parallel to polarity, leading to elongation without epidermal cracking. (I) Final state with reduced specified growth in epidermis, leads to a shorter cylinder and epidermal cracks. (J) Longitudinal section through (I), showing tissue tension in blue and compression in orange.

To further validate this hypothesis, we modelled the growth of a solid cylinder with epidermal and internal regions, in which cracks can form when tension exceeds a threshold value. The epidermal region in the model had four times the stiffness of the internal region, because average outer epidermal wall thickness was measured to be about four times greater (Fig. 4F, fig. S8). When specified growth in all tissue layers was equal, the cylinder elongated, and cracks did not form (Fig. 4H). Low specified growth in the epidermis led to reduced elongation, crack formation and tissue stresses (Fig 4, I to J), similar to what is observed in the BRZ-treated *qua2-1* mutant.

Our findings indicate that at early developmental stages, the epidermis of hypocotyls/stolons has a low specified growth rate that mechanically constrains the axial growth of internal tissue, generating tissue stresses. The most likely cause of the growth constraint is the thick outer epidermal cell wall, observed in both wild-type *Arabidopsis* and *Utricularia* at early developmental stages (Fig. 4F and fig. S8). Brassinosteroids can promote hypocotyl elongation within 6 h of application by increasing wall extensibility (16, 17), possibly via phosphorylation of plasma membrane H^+^-ATPase (18). Thus, if brassinosteroids preferentially increase extensibility of the thick outer cell wall, this would release internal tissue from mechanical constraint and promote axial growth. The cracks in hypocotyls of *qua2-1* mutants indicate that this release is not complete, and some tissue stresses remain. A constraining outer wall is also supported by the curved shape of epidermal cells where cracks form (Fig. 4D blue arrowheads). Wall-specific modification may be mediated by cell polarity factors that confer differences between cell faces (19-21)

This mode of brassinosteroid action suggests that the non-autonomous effect of epidermal brassinosteroid synthesis in promoting growth of dwarf mutants (4) may be mediated, at least in part, by release of tissue stresses. Mechanical interaction between cell layers have also been shown to play a role in formation of cracks in crocodile skin (22) and formation of intestinal villi (23). Here we reveal a likely mechanism whereby genes may influence such interactions. Thus, genetically controlled mechanical interactions between cell layers via tissue connectivity likely plays a fundamental role in coordinating morphogenesis in diverse systems.

## Acknowledgments

We would like to thank B. and P. Steward at The Fly Trap Plants and T. Bailey from the Carnivorous Plant Society for plants, seeds, and advice and Mateusz Madja for *qua2-1* seeds, Eva Wegel and Sergio Lopez JIC Bioimaging for help with light microscopy, Lionel Perkins and JIC Horticulture team for large scale *U. gibba* cultivation and Desmond Bradley for critical reading of the manuscript.

## Author contributions

RK-B, KL, JEB, MY, PB, BK, SC, JC, TX, BL, JF, YX, RS and CDW contributed biological experiments, data analysis and conceptualization, RK and EC computational modelling, RK software development, KL, CB, JS, MY and CDW development of *U. gibba* resources, RK-B and AW bioinformatic analysis, and EC supervision, funding acquisition and conceptualization.

## Data and materials availability

All data is available in the main text or the supplementary materials. Code is available at the following website: TBC.

## Competing interests

Authors declare that they have no competing interests.

Supplementary Materials

Materials and Methods

Figs. S1 to S9

References (24-33)

