## Supplementary materials for "Genetic control of cell layer interactions in plants via tissue mechanics"

### Materials and Methods

#### <u>*U. gibba* plant material and growth conditions</u>

##### Tissue culture

*Utricularia gibba* seeds of wild-type plants were purchased from Fly Trap Plants (Bergh Apton, UK). Plant material was grown in liquid ½ MS plant tissue culture media (0.22 % Murashige and Skoog Medium (MS) (Duchefa Biochemie M0233), 2.5 % sucrose, pH 5.8) and maintained in controlled environment room (CER) conditions at 23 ±1 °C, light at an intensity of 180 µmol/m$^2$/s, with a 16-h light/8-h dark photoperiod.

##### Glasshouse conditions

Plant material was grown in the glasshouse to induce flowering for seed collection. Plants were grown in containers containing a 2 cm layer of 1:1 peat:sand mix, topped up with reverse osmosis water.

##### Seed sterilisation

Seeds were washed for 5 minutes in 70 % ethanol, 0.1 % SDS, washed in sterile water and transferred to 4% bleach, 0.2 % triton 100 for 10 minutes, then washed 3 times with sterile water.

##### Seed germination

Seeds were sown in sterilin jars containing a layer of solid culture medium (0.22 % MS, 2.5 % sucrose, 0.3 % agar, pH 5.8) topped up with liquid MS culture medium containing 0.1 mM ethephon (Sigma C0143). To make ethephon containing media, a concentrated 2.5 M ethephon solution was made in a pH 3 buffer (41 mM disodium hydrogen phosphate, 79 mM citric acid) and diluted in liquid media to a final concentration of 0.1 mM. Seedlings were germinated at 23 °C in CER conditions (as above). Once seeds had germinated, seedlings were removed from ethephon containing media and grown in MS liquid media (as above).

### Mutagenesis of *U. gibba* tissue with EMS

*U. gibba* plants were grown in sterile culture prior to EMS treatment. Plant material was treated with 0.01 %, 0.05 %, 0.01 %, 0.15 %, 0.2 %, or 0.25 % EMS (ethyl methanesulfonate) diluted in 0.02 % tween 20 (Sigma-Aldrich, P9416). Tissue was incubated with the EMS solution while being continually agitated for 18 hours. Treated tissue was then passed through 10 x 20-minute washes in 0.02 % Tween, washed twice in water and incubated overnight in water and placed in the CER (as above). Tissue was divided into separate M1 plants in the glasshouse (approximately 5 cm of stolon). Flowering M1 plants had seed collected to produce M2 generation to identify segregating phenotypes of interest to take on to M3. For the family containing the dwarf and extreme-dwarf plants used in this study, phenotyping only separated plants into wild type, dwarf or extreme-dwarf plants, therefore subtleties in other segregating phenotypes may have been missed.

### Passaging *U. gibba*

Wild type, dwarf and extreme-dwarf plant material were treated in liquid culture using epibrassinolide (Sigma Aldrich, E1641) to provide exogenous brassinosteroid. Newly grown plant material was sub-cultured into fresh media containing fresh treatment every week to ensure exposure level. Treated plant cultures were maintained in CER conditions described above.

### Tracking stolons

2 cm length of dwarf plant stolon with an apex were isolated in sterilin jars for one week, imaged on a Leica M205C stereomicroscope with a Leica DFC495 camera (Leica, Milton Keynes, UK) at day 0 on a plate containing water, then again isolated in sterilin jars of liquid media containing the appropriate treatment and returned to CER . Individual stolons were imaged at day 7 then returned to fresh media containing the appropriate treatment for a week before being imaged at day 14. Individual images were stitched together in Adobe Photoshop and nodes labelled to identify internode 0 at day 0 and internodes nodes which had subsequently appeared in treatment at day 14 were labelled with negative numbers. Internode length measurements were made in ImageJ software (http://imagej.nih.gov/ij/).

### *A. thaliana* plant material and growth

### Tissue culture

*A. thaliana quasimodo2-1* plants were grown on plates containing MS media (0.441% Murashidge and Skoog including vitamins, 1% (w/v) glucose, 0.05% (w/v) MES, 1% Difco agar, pH to 5.7). Sterilised seeds were stratified in the dark at 4°C for 2 days, then exposed for light for 4 hours at 20°C in a controlled environment room before being wrapped in three layers of tin foil to ensure etiolation. To inhibit BR, 1 µM of BRZ was chosen that has been shown to replicate the phenotype of a BR biosynthesis mutant (*24*). BRZ was added to media of treated seeds which were subjected to the same conditions and untreated seeds.

### Seed sterilisation

Seeds were sterilised in 70% ethanol with 0.05% SDS for 5 minutes, followed by three washes in 100% ethanol. Seeds were air-dried on sterile filter paper before being plated (as above). If seeds were receiving hormonal treatment, then this was added to the media that the seeds would germinate on.

General methods

### Propidium iodide staining for confocal imaging

The propidium iodide staining protocol for whole-mount imaging (*25*) was followed to stain *U. gibba* with the following extra steps. After the final water wash, tissue was mounted onto glass slides with added Frame-Seal Incubation Chambers (BIO-RAD, SLF0601). A drop of ½ strength chloral hydrate solution was added to cover the tissue and samples were incubated over-night at room temperature. Excess chloral hydrate was removed and samples correctly spaced on the cover slip. Samples were mounted in Hoyer's solution and a slide placed on top to ensure samples were close to the coverslip for imaging.

For *qua2-1,* hypocotyls were placed in 0.25 mg/ml propidium iodide for 10 minutes, washed in water then placed on a glass slide with added Frame-Seal Incubation Chambers (BIO-RAD, SLF0601) plus water before imaging.

### Confocal imaging

Tissue samples were PI stained and mounted as described above. Imaging was performed using a x10 or x20 dry lens on a Zeiss 780 or 880 confocal microscope. 561 nm excitation was used, collected at 625-690 nm.

### Cell segmentation with MorphographX

Confocal Z-stacks were resized, brightness/contrast adjusted as required and converted to .tiff format with Image J (http://imagej.nih.gov/ij/). Stacks were loaded into MorphoGraphX open-source software (Software – MorphoGraphX) and processes described in (*26*) followed to create surface meshes and segment epidermal cells. Heatmaps for cell area, max and min cell length, anisotropy (cell max length/(cell max length + cell min length) and cell length parallel and perpendicular to a Bezier line drawn along the stolon axis were generated and data exported as .csv files and viewed in Microsoft Excel to generate charts.

#### Statistical Analysis

Statistical analysis was performed using R version 2022.07.1. to perform ANOVA with Tukey Test in Fig. 1 and S6 and *t*-test in Fig. 4.

### Light microscopy imaging

Live plant tissues were imaged in water using a Leica M205C stereomicroscope with Leica DFC495 camera. Plant phenotype measurements were taken using ImageJ software (http://imagej.nih.gov/ij/).

### Transmission electron microscopy

Stolons and hypocotyls were cut into small pieces and immediately placed in a solution of 2.5% (v/v) glutaraldehyde in 0.05M sodium cacodylate, pH 7.3 for fixation, and left overnight at room temperature. When samples were too thin for the smallest Leica EM TP baskets they were embedded in 2% (v/v) low gelling temperature agarose in water and plunged into ice. Once the agarose had set, $1mm^3$ blocks containing stolons or hypocotyls were cut out and placed in a solution of 2.5% (v/v) glutaraldehyde in 0.05M sodium cacodylate, pH 7.3 and left overnight to fix. The samples were loaded into a Leica EM TP embedding machine (Leica, Milton Keynes, UK) using the following protocol. The fixative was washed out by three successive 15-minute washes in 0.05M sodium cacodylate and post-fixed in 1% (w/v) $OsO_4$ in 0.05 M sodium cacodylate for one hour at room temperature. The osmium fixation was followed by three, 15-minute washes in distilled water before beginning the ethanol dehydration series (30%, 50%, 70%, 95% and two changes of 100% ethanol, each for an hour). Once dehydrated, samples were gradually infiltrated with LR White resin (London Resin Company, Reading, Berkshire) by successive changes of resin:ethanol mixes at room temperature (1:1 for 1hr, 2:1 for 1hr, 3:1 for 1hr, 100% resin for 1 hr then 100% resin for 16 hrs and a fresh change again for a further 8 hrs). Samples were transferred into gelatin capsules full of fresh LR White and placed at 60°C for 16 hrs to polymerize. The material was sectioned with a diamond knife using a Leica UC7 ultramicrotome (Leica, Milton Keynes, UK) and ultrathin sections of approximately 90nm were picked up on 200 mesh copper grids which had been formvar and carbon coated (EM resolutions, Sheffield, UK). The sections were stained with 2% (w/v) uranyl acetate for 1hr and 1% (w/v) lead citrate for 1 minute, washed in distilled water and air dried. The grids were viewed in a FEI Talos 200C transmission electron microscope (FEI UK Ltd, Cambridge, UK) at 200kV and imaged using a Gatan OneView 4K x 4K digital camera (Gatan, Cambridge, UK) to record DM4 files. For the visualisation of the material by light microscopy, semi-thin sections of 500nm were taken using a Leica Artos 3D ultramicrotome, stained with 0.5% (w/v) Toluidine blue and imaged on a Zeiss Axio Imager Z2.

### Freeze-fracture SEM

CryoSEM and cryofracture was carried out as described in (*27*) with the following modifications: (i) Iridium was used as the sputter coating target to a measured thickness of 3 nm and (ii) Imaging used the backscattered electron detector and a gun voltage of 25 kV and a probe current of 16 pA.

### Determination of endogenous BRs levels

Cathasterone analysis was performed at JIC, UK with extraction of purification as described in (*28*) and instrumental analysis as in (*29*). Typhasterol, 6-deoxocastasterone, and castasterone were detected using deuterium-labelled standards at Institute of Genetics and Developmental Biology, China. The quantification of endogenous BRs levels was performed based on the method reported previously with some simplifications in sample pretreatment (*30*). 200 milligrams of the sample powder was extracted with 90% aqueous methanol (MeOH) in an ultrasonic bath for 1 hour.

Simultaneously $D_3$-castasterone (CS), $D_3$-6-deoxocastasterone (6-deoxo-CS), and $D_3$-typhasterol (TY) were added to the extract as internal standards for BRs content measurement. After the MCX cartridge was activated and equilibrated with MeOH, water and 40% MeOH in sequence, the crude extracts reconstructed in 40% MeOH were loaded onto the cartridge. The MCX cartridge was washed with 40% MeOH, and then BRs eluted with MeOH. After drying with $N_2$ stream, the eluent was redissolved with ACN to be derivatized with 2-methoxypyridine-5-boronic acid (MPyBA) prior to UPLC-MS/MS analysis. BRs analysis was performed on a quadrupole linear ion trap hybrid MS (QTRAP 6500, AB SCIEX) equipped with an electrospray ionization source coupled with a UPLC (Waters) (*31*). As for CS, $D_3$-CS, 6-deoxo-CS, $D_3$-6-deoxo-CS, TY and $D_3$-TY, the MRM transition 582.4>178.1, 585.4>178.1, 568.4>178.1, 571.4>178.1, 566.4>548.3 and 569.4>548.3 was used for quantification.

### Sequence Analysis

Genomic DNA from 11 mutants, 13 individuals displaying a wild-type phenotype were sequenced at the Chinese Academy of Science, Beijing to a minimum of 35x coverage. Sequence data for 72 biologically identical progenitor samples were pooled together to identify novel mutations that were introduced to the mutant family. Libraries were prepared using a TruSeq Nano DNA kit and sequencing performed on an Illumina Hiseq X Ten to produce 150bp paired-end reads. Reads were mapped using Burrows-Wheeler Aligner (bwa-0.7.17) to the Chromium 10x reference created from the progenitor. Quality filtering was done by removing read overlaps using clipOverlap (bamutil-1.0.14) and PCR duplicates using MarkDuplicates (picard-1.134) with the following settings: REMOVE_DUPLICATES = true ASSUME_SORTED = true VALIDATION_STRINGENCY = SILENT MAX_FILE_HANDLES_FOR_READ_ENDS_MAP = 900. Variable sites were called using HaplotypeCaller (GATK-4.0.9.0) to identify variable sites. Initial filtering was performed (BCFtools-1.8) for biallelic sites with a minimum allelic count of 1 using the following command: bcftools view -m 2 -M 2 -O v -c 1:minor and tabulated using VariantsToTable (GATK-4.0.9.0) with -GT command to output genotypes for each individual at each variable site. The candidate gene was identified using the reference genome that has gene annotation (*32*). The coding sequence was extracted and annotated using Geneious (11.0.5).

Further genotyping was done using the KASP genotyping platform (LGC Genomics) using the VIC (5'-GAAGGTCGGAGTCAACGGATTAGGGGAGGAGCGGGCCTCGTGG-3') and FAM (5'-

GAAGGTGACCAAGTTCATGCTAGGGGAGGAGCGGGCCTCGTGA-3') fluorescent probes and a common reverse primer (5'- GTAGCTGCTTCTCGACGGCTCC-3').

### Supplementary Text

#### Modelling

All models were creating using GFtbox (https://coensoft.jic.ac.uk/software) and deposited at Github (link TBC).

#### Utricularia models (Fig. 2 J to V)

An initial mesh was created with an outer cylinder (epidermis) connected through six blades to an axial core (fig. S9A, B). For simulating the extreme-dwarf mutant, the initial mesh was shorter and had thicker internal blades and epidermis (fig. S3). The different regions of the mesh express different identity factors: EPIDERMIS, BLADES, and AXIS. Polarity initially ran from base to top of the cylinder and then deformed with the tissue. Specified growth was only parallel to the polarity and set to 4% per time unit for all regions in the wild-type model, and set to 0 only for EPIDERMIS in the dwarf mutant model. In all models the growth rate was set to zero in all regions near the top and bottom of the mesh. Residual strain (and thus stress) decayed by a factor of $e^{-0.5}$ in one time unit. At the start, the mesh was given a small random perturbation to the positions of all the vertexes, to break the symmetry and allow buckling. All models were run for 30 time units.

#### Limitations

The model assumes the growth constraint comes from slower specified growth of the epidermis, whereas in real tissue the constraint may come mainly from the outer wall of the epidermis. As only a small segment of stolon is modelled, the inner tissue bulges out from the top and bottom in dwarf mutants.

#### Arabidopsis models (Fig. 4 G to J)

An initial solid mesh was created as 6 concentric cylinders of finite elements (fig. S9C). The outer cylinder was assigned EPIDERMIS identity and the rest INNER identity (orange). The surface-half of the EPIDERMIS region was coloured blue and the internal half orange. The bulk modulus of the EPIDERMIS region was four times that of the inner region, reflecting the greater average wall thickness of the hypocotyl epidermis. Polarity was initially ran from base to top of the cylinder and then deformed with the tissue. Specified growth was only parallel to the polarity. For the control model (Fig. 4H), all regions had the same growth rate (4% per time step). For the BRZ-treatment model (Fig. 4I and J), specified growth rate of the EPIDERMIS region was set to 0, while that of the INNER region remained at 4%.

Cracks formed along boundaries of the finite elements when the residual tension exceeded a threshold level. To prevent the tissue all cracking at once, variation in weakness was generated in the mesh. A diffusible factor, WEAKNESS, was initially given an independent random value at every vertex. WEAKNESS set the strength of the tissue: the higher the value, the lower the tension at which it would crack. WEAKNESS was always in the range from 0 to the 'weakness' parameter. The distribution of WEAKNESS was smoothened by diffusion for the time specified by 'diffusiontime', before growth initiated. After each diffusion step, WEAKNESS was rescaled to the interval from 0 to 'weakness'. Setting WEAKNESS to 0 resulted in no cracks or internal twisting, even when specified growth was set to 0 in epidermis (fig. S7). After that time, its diffusivity was set to zero to freeze the pattern, and growth started. The parameter 'breakingstress' defined the residual stress required to make or extend a crack. The parameter 'strainretention' set the rate at which retained strain was retained. 0 = instant dissipation, 1 = no dissipation. 0.5 = decay by a factor of $e^{-0.5}$ in one time unit. For all models, 'strainretention' was set to 0.5.

### Limitations

The direction of the cracks is biased by the structure of the mesh, because the implementation of cracks only allows them to form along the boundaries of the finite elements. In real tissue, the pattern of cracks would follow the lines of weakness between cells, a feature not incorporated in the model. Also, finite elements cannot slide relative to each other, whereas in real tissue, cell sliding may allow cracks to open up more. As only a small segment of stolon is modelled, the inner tissue bulges out from the top and bottom in the BRZ model.

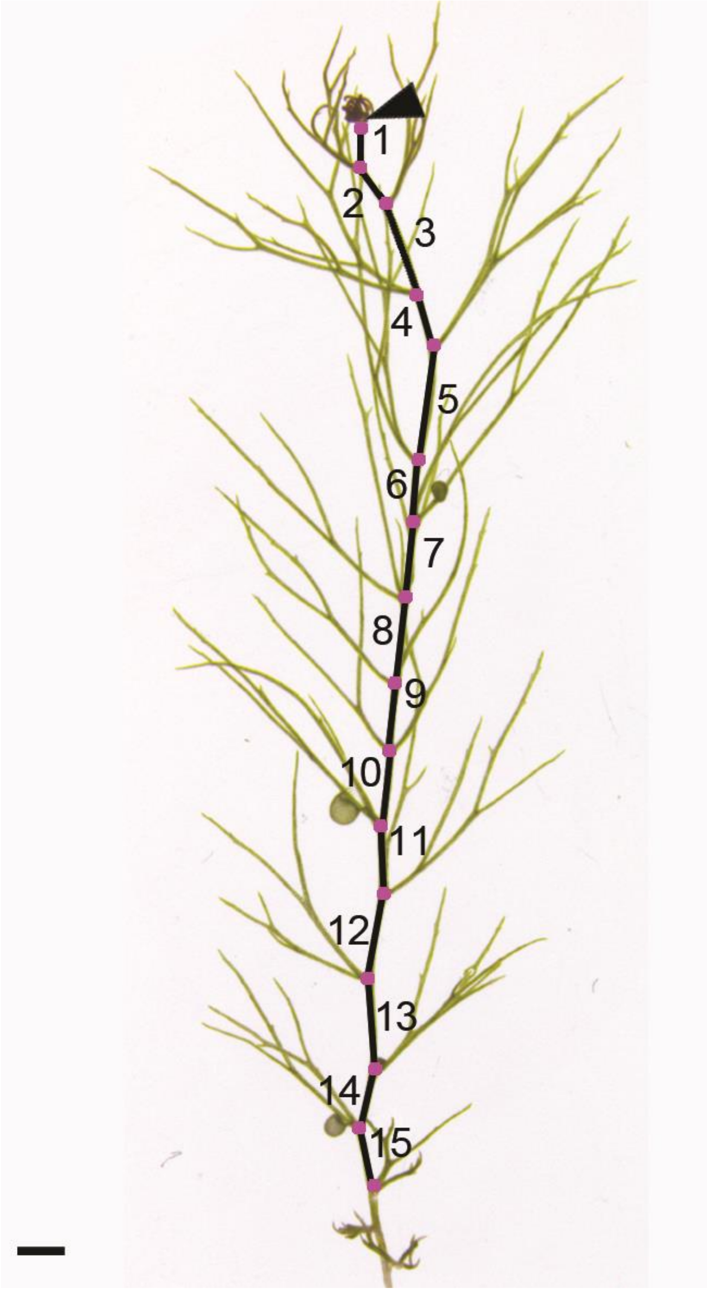


**Fig. S1. Internode numbers increase with stolon maturity**

Black arrowhead indicates first fully emerged leaf from apex, used to identify the beginning of internode 1. Nodes shown with magenta dots. Scale bar 1 mm.

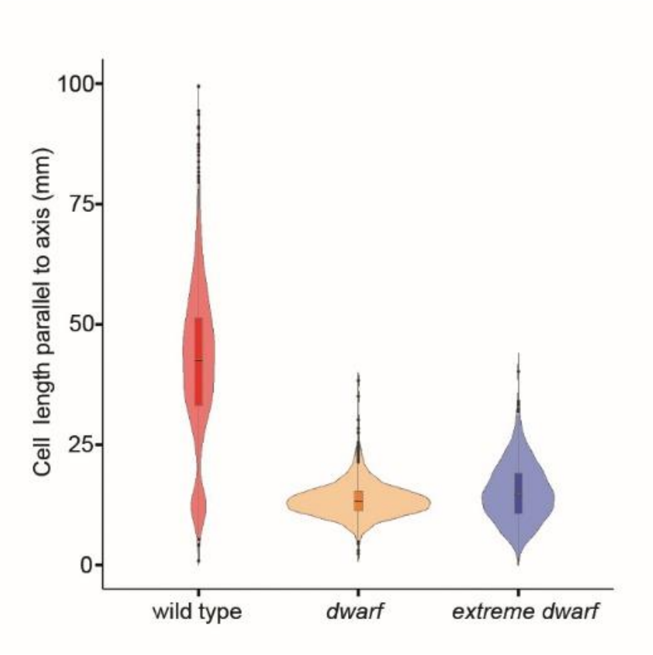


**Fig. S2. Cell length parallel to stolon axis**

Violin plots of cell length parallel to stolon axis of mature stolons of wild type (n=1817 cells from 8 plants), dwarf (n=2289 cells from 5 plants) and extreme-dwarf (n=1494 cells from 4 plants). Both mutants have significantly lower values than wild type ($p < 0.001$). Axial cell length was measured in relation to a manually placed Bezier line which ran the length of the stolon.

Mean cell length parallel to the axis is 41.5 µm for wild type, 13.5 µm for dwarf, and 15.1 µm for extreme-dwarf, showing axial cell length in dwarf mutants is reduced by a factor of 3, and extreme-dwarf a factor of 2.7. By comparison, mean internode length was 3.07 mm for wild type, 0.72 mm for dwarf and 0.29 mm for extreme-dwarf (Fig. 2D), indicating dwarf internode length is reduced by a factor of 4.25, and extreme-dwarf by a factor of 10.5. These findings suggest that about 70% (3/4.25) of the reduction of dwarf internode length, and 26% (2.7/10.5) of the reduction of extreme-dwarf internode length were caused by reduced cell length, with the remaining reduction caused by reduced cell number.

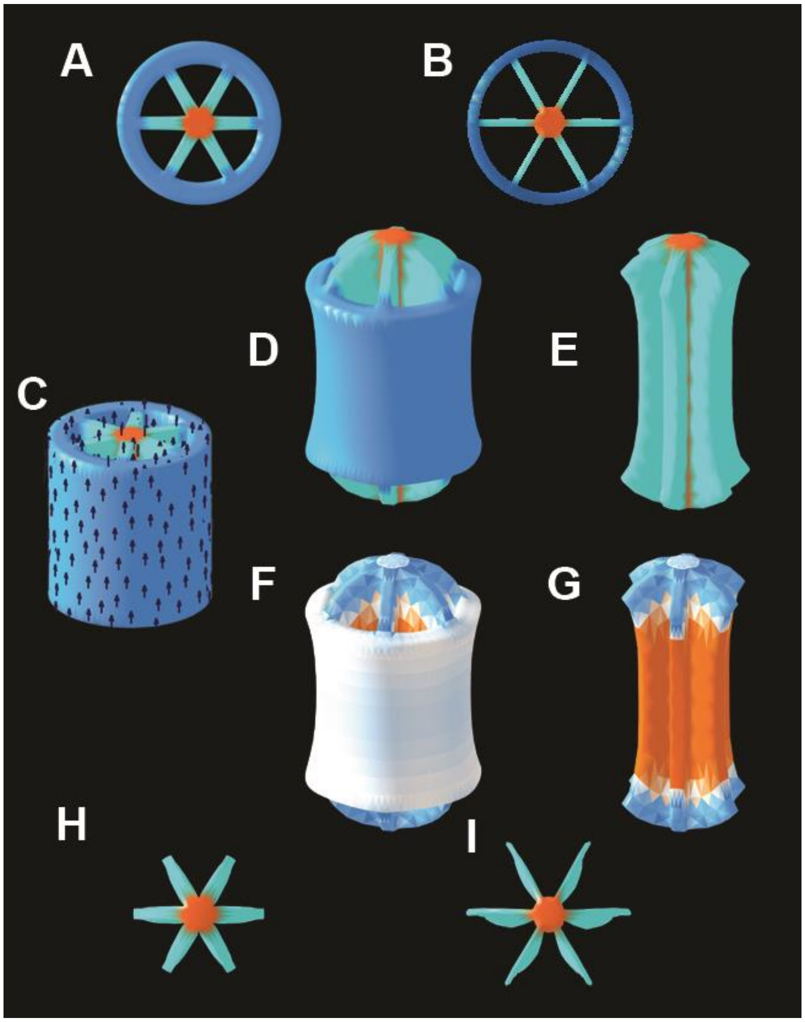


**Fig. S3. Model of extreme-dwarf when air spaces are smaller**

(**A** and **B**) Top-down views of initial state with small air spaces (A), compared to the larger air spaces (B) used in the models of Fig. 2. Epidermis (blue), blades (cyan) and axial core (orange). (**C**) Side view of extreme-dwarf model initial state with smaller air spaces. Polarity shown as black arrows. (**D**) Final state when epidermis has low specified growth rate. (**E**) as (D) with epidermis removed. (**F** and **G**) as (D and F) showing tissue compression in orange and tension in blue. (**H**) Top-down view of (E). (**I**) Top-down view of Fig. 2T for comparison.

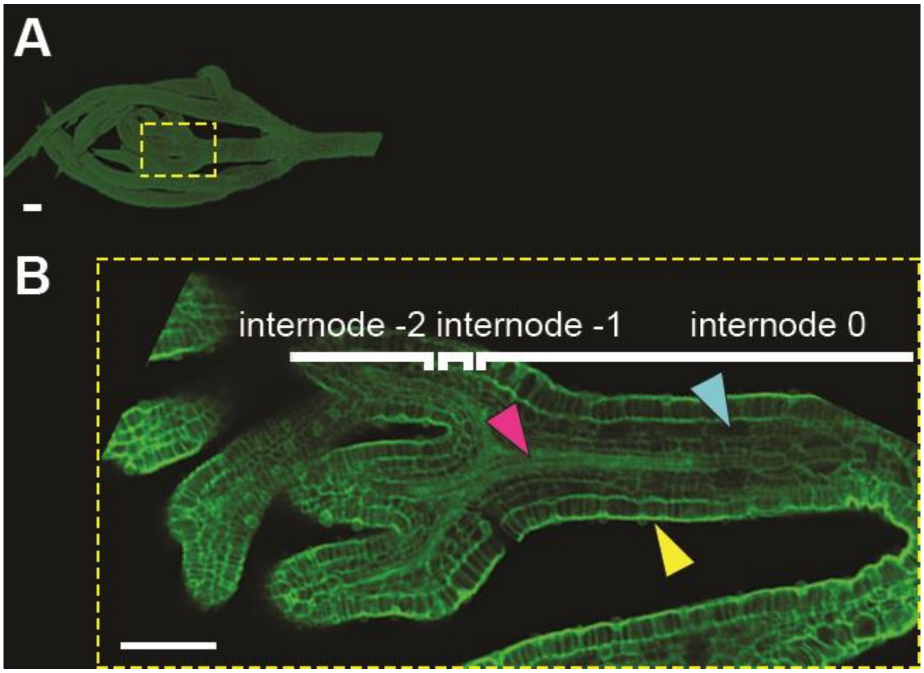


**Fig. S4. Wild type inner tissue organization at early stages**

(**A**) Confocal scan of early wild type internodes. (**B**) Enlargement and z-slice of (A). Vasculature (magenta arrowhead), air spaces (cyan arrowhead) and epidermis (yellow arrowhead). Scale bars 100 µm.

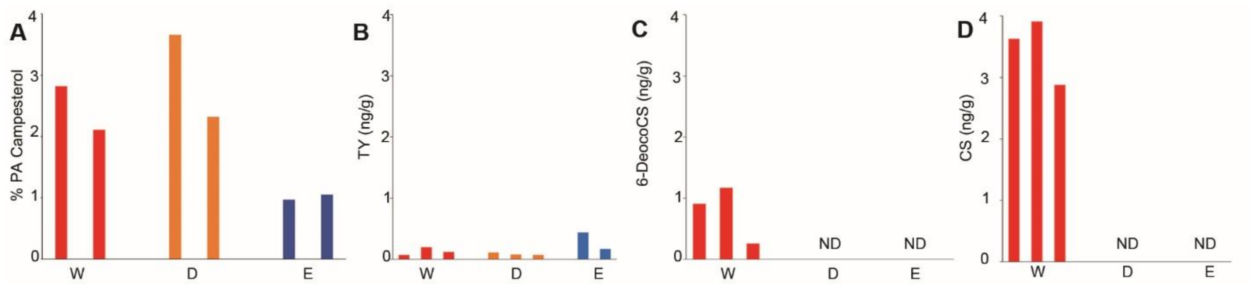


### Fig. S5. Brassinosteroid precursors in dwarf plants of *U. gibba*

(**A**) Level of Campesterol, a BR precursor upstream of the *dwarf4* block in *Arabidopsis*. As no internal standard was available for Campesterol, the level of Campesterol was calculated by the percentage of the chromatogram area occupied by its peak. Replicates shown for different individuals. (**B-D**) Levels of precursors downstream of the *dwarf4* block. (B) Typhasterol (TY), (C) 6-Deoxocastasterone (6-DeoxoCS) and (D) Castasterone (CS) in ng/g of tissue. TY is present in all samples but at low levels and therefore could be attributed to noise. Precursors are shown from left to right in their position on the biosynthetic pathway (*33*). Wild type (W, red), dwarf (D, orange) and extreme-dwarf (E, blue). ND = not detected.

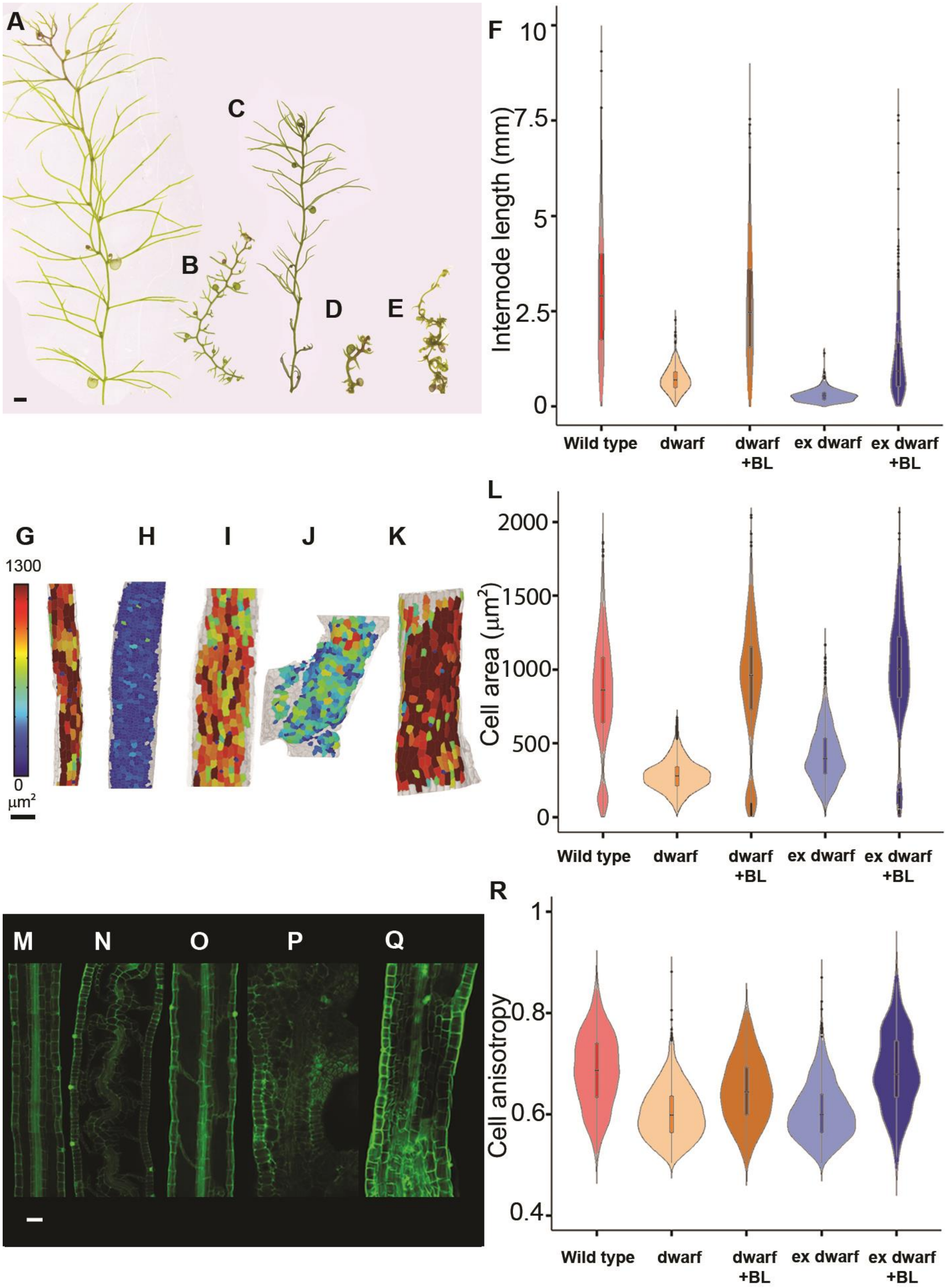
A
B
C
D
E
F
Internode length (mm)
10
7.5
5
2.5
0
Wild type
dwarf
dwarf +BL
ex dwarf
ex dwarf +BL
G
H
I
J
K
1300
0
µm²
L
Cell area (µm²)
2000
1500
1000
500
0
Wild type
dwarf
dwarf +BL
ex dwarf
ex dwarf +BL
M
N
O
P
Q
R
Cell anisotropy
1
0.8
0.6
0.4
Wild type
dwarf
dwarf +BL
ex dwarf
ex dwarf +BL

### Fig. S6. Treating with brassinolide rescues dwarf, and partially rescues extreme-dwarf.

Phenotypes and data from Figs 1 and 2 shown for comparison. (A to E) Whole-plant phenotypes. (A) Wild type. (B) Dwarf (C) Dwarf + 0.01 uM epiBL. (D) Extreme-dwarf. (E) Extreme-dwarf + 0.01 uM epiBL. Scale bar 1 mm. (**F**) Violin plots of mature internode lengths of wild-type (n=10, as in Fig. 1D), dwarf (n=10, as in Fig. 1D) dwarf + 0.01 uM epiBL (n=19), extreme-dwarf (n=13, as in Fig. 1D) and extreme-dwarf + 0.01 uM epiBL (n=15). Plants were grown in continuous culture. Block indicates interquartile range and horizontal line the mean. Both treated mutants had greater lengths than untreated ($p < 0.001$). Dwarf + 0.01 uM epiBL was not significantly different from wild type ($p = 0.768$). (**G** to **K**) Heat maps of cell area in mature stolons. (G) Wild type. (H) Dwarf. (I) Dwarf + 0.01 uM epiBL. (J) Extreme-dwarf. (K) Extreme-dwarf + 0.01 uM epiBL. Scale bar 100mm. (**L** and **R**) Violin plots of cell area (L) and cell anisotropy (cell max length/(cell max length + cell min length)) (R) of mature stolons of wild type (n=1817 cells from 8 plants, as in Fig.1 I and J), dwarf (n=2289 cells from 5 plants, as in Fig.1 I and J), dwarf + 0.01 uM epiBL (n=721 cells from 3 plants), extreme-dwarf (n=1494 cells from 4 plants, as in Fig.1 I and J) and extreme-dwarf + 0.01 uM epiBL (n =327 cells from 2 plants). Both treated mutants had increased lengths compared untreated ($p < 0.001$, L; $p < 0.01$, R). Both dwarf + 0.01 uM epiBL and ex dwarf + 0.01 uM epiBL were not significantly different to wild type cell area ($p = 0.99$ for dwarf+BL and $p=0.30$ for ex+BL) and cell anisotropy ($p = 0.34$ for dwarf+BL and $p=0.99$ for ex+BL). (**M** to **Q**) Longitudinal confocal sections. (M) Wild type. (N) Dwarf. (O) Dwarf + 0.01 uM epiBL. (P) Extreme-dwarf. (Q) Extreme-dwarf + 0.01 uM epiBL. Scale bar 50 µm.

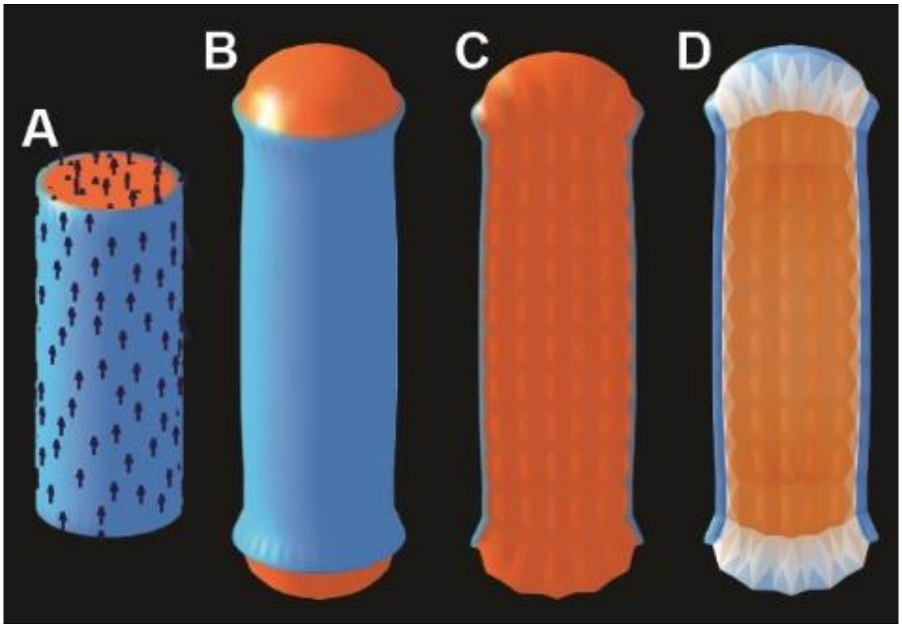


**Fig. S7. Reduced epidermal specified growth of a cylinder with solid internal tissue and no cracking**

(**A**) Initial state of model, as in Fig. 4G. (**B**) Final state with reduced specified growth in epidermis leads to a shorter cylinder. (**C**) Longitudinal section through (B), showing internal tissue does not twist. (**D**) Longitudinal section showing internal tissue tension in blue and compression in orange.

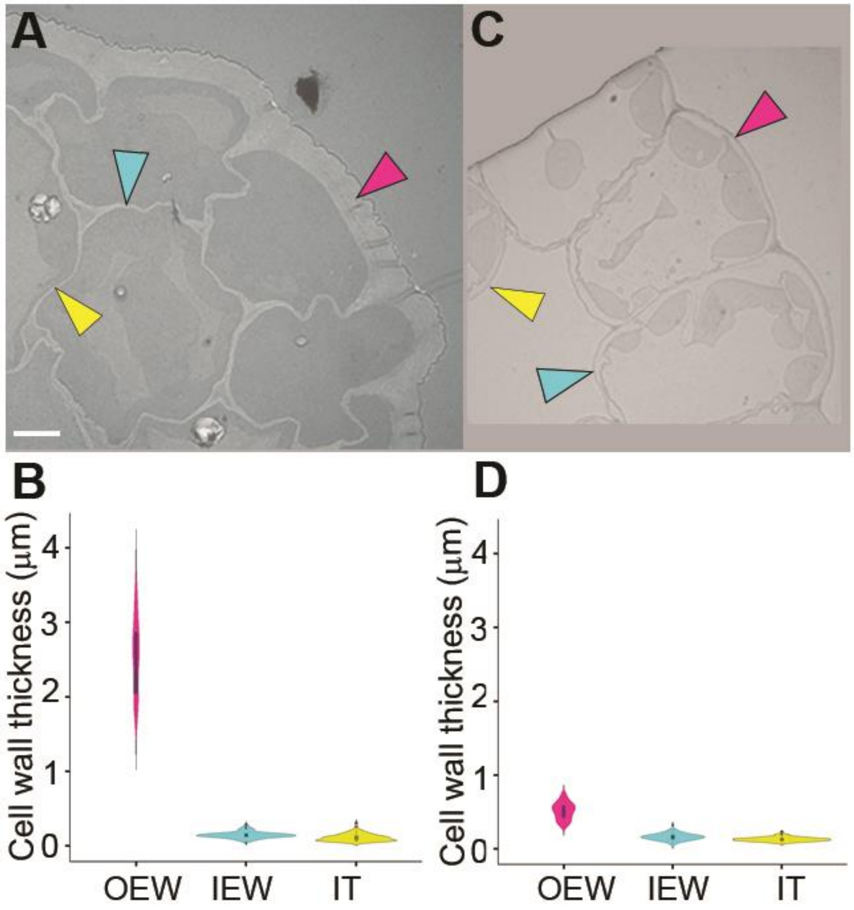


**Fig. S8. Outer epidermal walls are thicker than internal cells in both *Arabidopsis* and *Utricularia* at early stages**

(**A**) Transmission electron micrograph of transverse section through a wild-type Arabidopsis hypocotyl after 4 days growth, showing thicker outer epidermal cell wall. Outer epidermal wall (OEW, magenta arrowhead), inner epidermal wall (IEW, cyan arrowheads) and inner tissue cell walls (IT, yellow arrowheads). Scale bar 5 µm. (**B**) Violin plots of average cell wall thickness of OEW ($\bar{x}$ = 2.56 µm, n = 76 from 3 plants), IEW ($\bar{x}$ = 0.15 µm, n = 126 from 3 plants) and IT ($\bar{x}$ = 0.11 µm, n = 90 from 3 plants). Block indicates interquartile range and horizontal line the mean. OEW is greater than both inner epidermal walls and inner tissue walls ($p < 0.001$). Average thickness of epidermal cell walls was calculated from transverse sections by first measuring the mean fraction, f, of the epidermal cell perimeter occupied by the OEW (f = 0.303). Average epidermal cell wall thickness = f x average OEW thickness + (1-f) x average IT wall thickness = 0.852 µm. The ratio of this value to average IT wall thickness (0.11 µm) was 7.749. (**C**) Transmission electron micrograph of wild-type *U. gibba* internode 1. Scale bar 5 µm. (**D**) Violin plots of average cell wall thickness of OEW ($\bar{x} = 0.52$ µm, $n = 103$ from 3 plants), IEW ($\bar{x} = 0.16$ µm, $n = 84$ from 3 plants) and IT ($\bar{x} = 0.13$ µm, $n = 109$ from 3 plants). Block indicates interquartile range and horizontal line the mean. OEW is greater than both inner epidermal walls and inner tissue walls ($p < 0.001$).

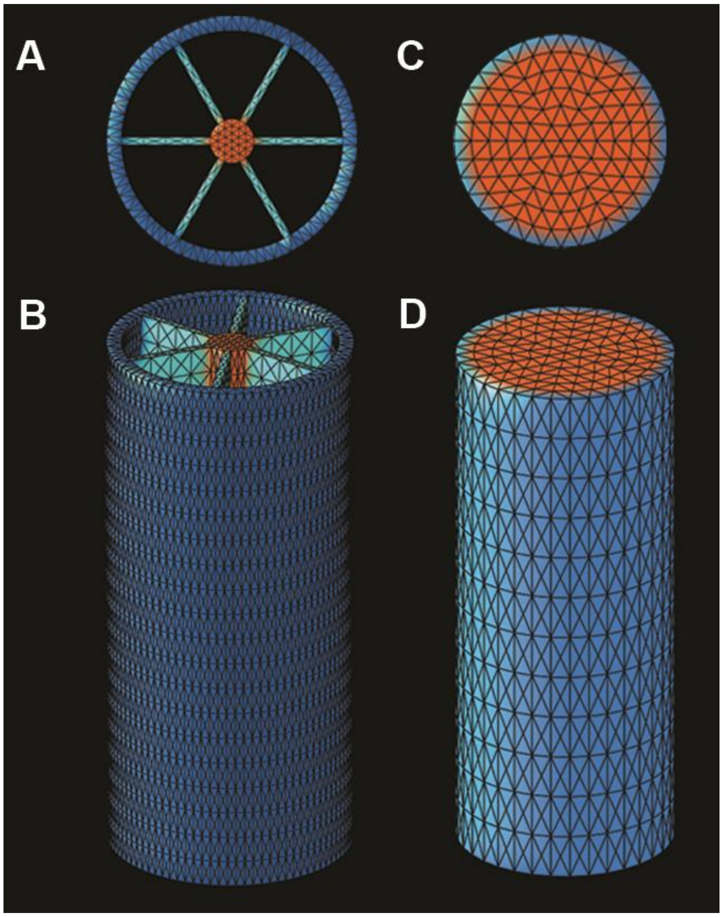


### Fig. S9. Finite element meshes of *Utricularia* and *Arabidopsis* models

(**A** and **B**) Initial mesh for *Utricularia* model was created with an outer cylinder (epidermis, blue), an axial core (orange) and six connective blades (cyan). (A) Top-down view. (B) Side view. (**C**) A solid mesh cylinder consisting of six concentric rings of finite elements. The outermost ring was assigned epidermal identity (outer surface blue, inner orange) and the other rings were treated as inner tissues (orange).
